# Invasive ants fed spinosad collectively recruit to known food faster yet individually abandon food earlier

**DOI:** 10.1101/2024.06.20.599949

**Authors:** Henrique Galante, Moritz Forster, Cosmina Werneke, Tomer J. Czaczkes

## Abstract

Current management strategies applied to invasive ants rely on slow-acting insecticides which aim to delay the ant’s ability to detect the poison until its effects are noticeable. Despite this, most control efforts are unsuccessful, likely due to bait abandonment and insufficient sustained consumption. Conditioned taste aversion, a learned avoidance of a particular taste, is a crucial survival mechanism which prevents animals from repeatedly ingesting toxic substances. However, whether ants are capable of this delayed association between food taste and subsequent illness remains largely unexplored. Here, we exposed colonies of the highly invasive Argentine ant, *Linepithema humile*, to a sublethal dose of the slow-acting insecticide spinosad. We combined measurements of individual-level feeding patterns with quantification of collective preferences and foraging dynamics to investigate the potential effects of the toxicant on behaviour. Collectively, ants preferred an odour associated with a previously experienced food, even if this contained spinosad, over a novel one. However, at the individual-level, previous exposure to spinosad resulted in reduced food consumption, as a consequence of earlier food abandonment. Moreover, while control-treated colonies recruited slower to a food source which tasted like a previously experienced one, spinosad-exposed colonies recruited equally fast to both novel and familiar foods. Although it appears that ants are unable to develop a conditioned taste aversion to sublethal doses of spinosad, ingestion of even small amounts of the toxicant strongly influences foraging behaviour. Understanding the subtle effects of slow-acting pesticides on ant cognition and behaviour can ultimately inspire the development of more efficient control methodologies.

## Introduction

Argentine ants are among the most widespread and destructive invaders globally, responsible for significant ecological and economic damage (Silverman & Brightwell, 2008; Angulo *et al*., 2022, 2024). Current pest control efforts often rely on chemical baiting. However, no successful eradication of an entire population of *L. humile* using slow-acting toxic hydrogel beads has been reported. This is thought to be due to low sustained bait uptake and the active abandonment of foraging trails to baits (Silverman & Brightwell, 2008; Zanola, Czaczkes & Josens, 2024). We hypothesise that this could, in part, be due to ants becoming averse to the taste of the toxicants in the baits. Spinosad, a slow-acting neuroactive toxicant, acts by stimulating nicotinic acetylcholine and GABA receptors, inducing rapid nervous system excitation, resulting in insect paralysis and death (Salgado, 1998; Biondi *et al*., 2012). Due to its low toxicity to mammals and fish, it is a promising eco-friendly insecticide for invasive ant management (Bacci *et al*., 2016; Khan, 2018). However, it is unclear if ants are able to detect its presence in the baits and if they become averse to it once its negative post-ingestion effects begin.

Learning allows animals to adapt to environmental changes throughout their lives. Associative learning, in particular, links an unconditional stimulus (a stimulus that causes a specific response without prior learning) with a conditional one (a stimulus that can be perceived but does not result in that specific response). Once these stimuli are associated, detecting the conditional stimulus triggers a response similar to that caused by the unconditional stimulus. This type of learning often requires multiple pairings of the unconditional and conditional stimuli, which must occur in close temporal proximity (Pavlov, 1927; Rescorla & Wagner, 1972; Dickinson, 2012). However, conditioned taste aversion (CTA), a form of classical conditioning, is an exception to these requirements.

CTA is a learned avoidance of a particular taste, developed when an initially neutral taste is associated with post-ingestion malaise, a general feeling of discomfort often linked to illness. This mechanism is crucial for survival, as it prevents animals from repeatedly ingesting toxic substances. In mammals, where CTA has been extensively studied, it is characterized by four main features: 1) a single conditioning trial is sufficient for forming long-lasting aversion (Steinert, Infurna & Spear, 1980; Rosas & Bouton, 1996); 2) conditioning can occur with a long delay of up to several hours between the conditional and unconditional stimuli (Nachman, 1970); 3) it is more easily achieved with a novel taste rather than familiar ones (Bernstein, 1999); and 4) illness is more easily associated with food taste than with other sensory cues, unless odour is compounded with taste (Garcia & Koelling, 1966; Palmerino, Rusiniak & Garcia, 1980).

Animals can learn to avoid odours associated with toxic food through both pre-ingestion and post-ingestion mechanisms. In honeybees, these mechanisms are mediated by different neurotransmitters: dopamine pathways primarily mediate direct aversion based on food unpalatability, while serotonin is involved in CTA signalling (Wright *et al*., 2010; Wright, 2011; Lai *et al*., 2020). Harnessed honeybees have been shown to develop aversions to bitter, toxic substances due to the physiological consequences of ingestion, with the malaise generated by these substances leading to a decreased response to odours which were previously appetitive (Ayestaran, Giurfa & Sanchez, 2010). On the other hand, honeybees did not develop a conditioned taste aversion to ethanol (Varnon *et al*., 2018). This suggests, that although honeybees are capable of CTA, not all compounds which likely cause post-ingestion malaise can be successfully associated with their negative effects.

In crickets, CTA is similar to that in mammals in that a single trial pairing is sufficient to achieve long-term memory retention. However, it differs in that the interval between food ingestion and toxin exposure has to be relatively short, under one hour (Lyu & Mizunami, 2022). Snails also exhibit CTA, which remains intact with repeated presentations of the conditioned stimulus and is selective for novel tastes (Nakai *et al*., 2020). Additionally, other insects such as moth larvae (Dethier, 1980), grasshoppers (Bernays & Lee, 1988; Simões, Ott & Niven, 2012), and fruit flies (Babin *et al*., 2014; Kobler *et al*., 2020) have been shown to form CTA.

Ants, known for their strong associative learning and reliance on multimodal cues, are incredibly fast learners (Knaden & Graham, 2016; Arenas & Roces, 2018; Oberhauser *et al*., 2019; Piqueret, Sandoz & d’Ettorre, 2019; Czaczkes & Kumar, 2020). Specifically, Argentine ants, *Linepithema humile*, form long-lasting olfactory associations often after a single experience (Rossi *et al*., 2020; Wagner *et al*., 2023; Galante & Czaczkes, 2024). Moreover, while most ant research focuses on appetitive learning, ants can also form associations through aversive learning, such as odour-heat associations (Desmedt *et al*., 2017) or the avoidance of quinine, a bitter substance (Wenig, Bach & Czaczkes, 2021). However, whether they are capable of conditioned taste aversion remains largely unexplored.

In this study, we provided *L. humile* colonies with spinosad-laced sucrose solutions to determine the extent to which they can develop a conditioned taste aversion for this toxicant. Using Y-mazes, we compared collective preferences for two odours before and after colonies were fed flavoured spinosad-laced sucrose solutions. Additionally, we quantified individual feeding patterns and collective foraging dynamics in order to understand if spinosad is detected and aversive to the individuals, but also how these effects translate to collective decision making and foraging. Ultimately, understanding how ants react to toxicant-laced baits and how their foraging behaviour changes in response to this is crucial for the design of effective control strategies. If ants become averse to slow-acting toxic baits, this could drastically reduce the effectiveness of control attempts, and explain the general lack of success of current methodology.

## Materials and Methods

To determine whether ants can form a conditioned taste aversion to sublethal doses of spinosad and to assess the influence of this slow-acting toxicant on collective foraging behaviour we combined three types of experiments. Firstly, we tested multiple individuals from each colony for their innate preference for two odours, apple and strawberry. We then fed 10% of each colony one of four treatments: apple-flavoured sucrose, strawberry-flavoured sucrose, apple-flavoured spinosad-laced sucrose, or strawberry-flavoured spinosad-laced sucrose. Approximately 24 hours later, we again tested multiple individuals from each colony for their post-ingestion preference for the same odours. We then quantified the amount and rate at which individuals which were not directly fed the treatment drank on either a food source with a novel taste-odour or one matching that of the treatment. Finally, we provided entire colonies two food sources - one with a novel taste-odour and one matching the previously experienced odour - and quantified their collective foraging dynamics.

### Colony maintenance

*Linepithema humile* (Mayr, 1868) were collected from Portugal (Alcácer do Sal and Proença-a-Nova) and Spain (Girona) between May 2021 and November 2022. Ants were split into colony fragments, containing three or more queens and 200-1000 workers, kept in non-airtight plastic boxes (32.5 x 22.2 x 11.4 cm) with a plaster of Paris floor and PTFE coated walls. 15mL red transparent plastic tubes, partly filled with water, plugged with cotton, were provided as nests. Ants were maintained on a 12:12 light:dark cycle at room temperature (21-26 °C) with *ad libitum* access to water. Between experiments, ants were fed *ad libitum* 0.5M sucrose solution and *Drosophila melanogaster* twice a week. From these established colony fragments, standardised experimental colonies (henceforth colonies) composed of 500 workers (nurses and foragers, randomly chosen), two queens and as little brood as possible were created. Colonies were then allowed to acclimate for one week and provided with *ad libitum* 0.5M sucrose solution and water. After one week, colonies were deprived of carbohydrates for four days prior to testing, ensuring high foraging motivation (Figure 1A). Experiments were conducted between November and December 2022 using 19 colonies.

**Figure 1.**
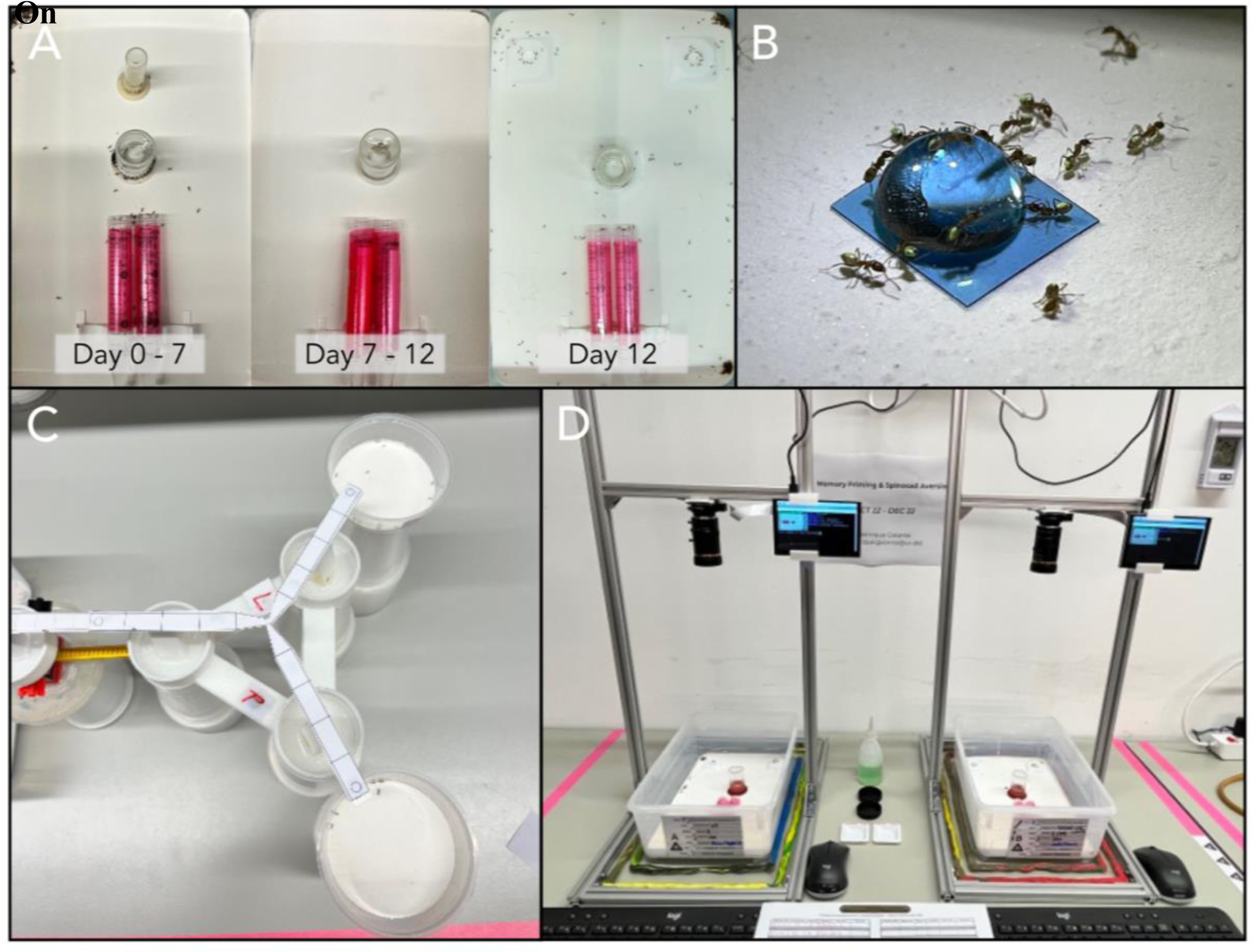
Overview of experimental setups and timeline of data collection. **(A)** Standardised experimental colonies (500 workers, 2 queens) were maintained from day 0 to day 7 with *ad libitum* access to both 0.5M sucrose solution and water. On day 7, colonies were deprived of carbohydrates for four days. On day 11, colonies were tested for their innate preference for two odours and then fed one of four possible treatments. On day 12, colonies were assessed for their post-ingestion preference, the feeding patterns of approximately 30 workers were measured, and collective foraging dynamics were recorded while colonies foraged on two food sources. **(B)** Polyacrylamide hydrogel bead containing one of four possible treatments, fed to 50 individuals which were marked with acrylic paint during ingestion. **(C)** Y-maze preference test setup connected to a colony, allowing individuals to make a choice between two odours. **(D)** Raspberry Pi HQ camera systems used to record collective foraging dynamics.

### Chemicals and solutions

Spintor (44% w/w spinosad), a commercial insecticide, was obtained from Corteva Agriscience (Brussels, Belgium). 1M sucrose solutions (Südzucker AG, Mannheim, Germany) mixed with 0.25ppm of spinosad were used as treatments. This concentration was chosen based on previous mortality reports suggesting it as a potential LD_50_ of *L. humile* kept in groups of 10 individuals (Galante et. al. In Prep.). Identical 1M sucrose solutions were used as controls. All solutions were mixed with 2μL/mL of either apple or strawberry food flavouring (Seeger, Springe, Germany). Scented paper overlays, used during the Y-maze preference test, were stored for at least one week prior to the experiments in airtight plastic boxes (19.4 × 13.8 x 6.6 cm) containing a glass petri dish with 0.5mL of either apple or strawberry food flavouring. Polyacrylamide hydrogel beads, used as food sources in the collective foraging test, were prepared one day prior to use. This was done by mixing 10mL of one of the four treatment solutions with approximately 80mg beads (around 8 beads) in a 15mL plastic tube. The beads were left to absorb the solution for about 24 hours at 6°C. For ease of use, the beads were cut in half before being placed on a platform, ensuring they lay flat.

### Y-maze innate preference test

After depriving the colonies of carbohydrates for four days, approximately 50 individuals were given access to a drawbridge connected to a Y-maze (Figure 1B). The Y-maze consisted of three 10cm long, 1cm wide arms, tapering to 2mm at the bifurcation (Czaczkes, 2018). One arm of the Y-maze had a removable paper overlay scented with apple flavour, while the other arm had one scented with strawberry flavour, both novel stimuli for the ants. This setup allowed the ants to choose a side without any prior learning, thus measuring their innate preference for the two scents. To control for potential side biases due to brain lateralization, we tested the 50 individuals in two sequential trials with 25 ants each. In the second trial, the arm of the Y-maze in which the paper overlays were placed was swapped. Once an ant chose a side, it was removed from the maze and kept separately until all 50 ants had made their choices, after which they were returned to their original colony. In total 918 individuals from 19 colonies were used.

### Treatment administration

Following the initial preference test, 50 workers from each colony were fed in small groups of three to five ants outside their nest with one of the four treatment solutions via polyacrylamide hydrogel beads (Figure 1C). These ants were marked with acrylic paint and kept in a separate box (19.4 × 13.8 x 6.6 cm) with *ad libitum* access to the treatment solution until all individuals were fed. This number, representing around 10% of the colony, corresponds to the expected proportion of foraging individuals in a colony, which usually falls within 10-20% of the workers in a colony (Lewis, Pollard & Dibley, 1974; Bruin, Röst & Draisma, 1977; Porter & Jorgensen, 1981; Hölldobler & Wilson, 2009). This method ensured a standardised amount of food was ingested by each colony regardless of treatment. Argentine ants visibly expand while feeding, which allowed us to confirm all treated individuals ingested the solutions provided. This approach also demonstrates that the solutions were palatable and thus it is unlikely there was any pre-ingestion aversion. Once fully fed, treated workers were returned to their respective colonies and allowed to share food and information through trophallaxis.

### Y-maze post-ingestion preference test

Approximately 24 hours later, the preference test was repeated with another batch of 50 ants, again divided into two groups of 25. These ants had presumably been exposed to the treatnebt solutions through trophallaxis. Ants were allowed onto a Y-maze with scented paper overlays, and their preference for the odour of the flavour with which they were treated was measured and later compared to their innate preference for that scent. After testing, ants were returned to their colony. In total 841 individuals from 19 colonies were recorded.

### Quantification of feeding patterns

Following the final Y-maze test, around 30 unmarked workers from each colony were removed and their feeding patterns, both total volume ingested and consumption rate, were quantified using 3D body reconstruction following Galante, Czaczkes & De Agrò (2024). Half of the individuals tested were fed a sucrose apple solution and the other half a sucrose strawberry solution. The setup consisted of a resin 3D-printed platform where the test solution was placed, attached to a Raspberry Pi HQ camera system and tracked using DeepLabCut version 2.3.8 (Mathis *et al*., 2018; Nath *et al*., 2019). The same DeepLabCut networks trained in Galante, Czaczkes & De Agrò (2024) were used, but the location of the cameras was adjusted and thus a new calibration performed (1px = 1.05 ± 0.20mm, N = 505). Each ant was recorded for its entire feeding event and removed from the platform once full, yet not returned to its colony of origin. In total 516 feeding events were recorded, 11 of which were excluded due to unsuccessful tracking and 102 manually removed due to the ants being in an unsuitable position for 3D reconstruction. Additionally, 49 recordings were removed due to a poor regression fit of volume over time (R^2^ < 80%) and four due to implausibly large crop load values. In total, 350 feeding events were analysed.

### Collective foraging dynamics

Finally, each colony was given access to two food sources, a sucrose apple solution and a sucrose strawberry solution. These solutions were provided as polyacrylamide hydrogel beads and placed on plastic platforms. Empty platforms were added when colony starvation began, ensuring ants were familiar with them. These were positioned approximately 3cm from each side, 4cm from the back, 13cm apart from each other, and 15cm from the nest entrance. To account for discovery time, which could significantly impact recruitment to one food source over the other (Sumpter & Beekman, 2003), one ant was placed at each food source after both beads were added. Colonies were recorded for four hours at one frame per second while foraging (Figure 1D). In total, 19 colonies were recorded.

### Automated quantification of ant count

ImageJ/Fiji (Schindelin *et al*., 2012; Schneider, Rasband & Eliceiri, 2012) was used to automatically count the number of ants at each feeder during the collective foraging experiment. An ImageJ macro was developed and applied to each frame (videos recorded at 1fps). First, a gaussian blur filter (radius = 0.4 pixels) was applied to the red colour channel of each frame. Following this, background subtraction was performed using a rolling ball method (radius = 2 pixels), the image’s contrast was enhanced (saturated pixels = 10%) and the Yen’s thresholding method was used (Yen, Chang & Chang, 1995). Watershed separation was applied to separate connected individuals. Pixels of intensity lower than 150 were removed, and colony-specific regions of interest (ROI) were set as circles (Ø 80 pixels) around each of the two hydrogel bead feeders. Using particle analysis, all between 15 and 200 pixels in size and between 0.3 and 1.0 in circularity within each of the ROI’s were counted. The output of this was a file containing the number of ants at each food source for every frame of each video. To validate the method, manual counts were performed blindly on ten images of each colony (N = 190). Selected images were evenly distributed across the duration of the videos. Manual ant count and ImageJ derived ant count had an almost perfect agreement of 98.6% (Spearman’s rank correlation coefficient, N = 380). Thus, ImageJ was as reliable as the manual counts, and therefore considered to be accurate.

### Quantification of collective foraging dynamics

Collective foraging followed a general pattern for all colonies and feeders. Initially, ant count rapidly increased (recruitment phase), reaching a maximum value which remained stable for a long period of time (plateau phase), and eventually began gradually decreasing (decline phase). For each of the two food sources in each colony, the end of the recruitment phase was taken as the point at which the maximum ant count was first reached. Similarly, the end of the plateau phase was taken as the point at which the maximum ant count was last reached. For all analysis, the decline phase was excluded, as it was most likely a result of the visible desiccation of the polyacrylamide beads (Cabrera *et al*., 2021), and thus food depletion. Moreover, the length and rate of both the plateau and the decline phase did not differ across treatments (see Statistical Analysis).

To quantify the rate at which colonies recruited to each food source, and the maximum ant count reached at each of them, a non-linear least-squares Michaelis-Menten model (Michaelis & Menten, 1913) was fit to the ant count over time data. Originally, the model describes biochemical kinetics and relates biochemical reaction rate to substrate concentration. However, the same formula can be applied to collective foraging dynamics, where the ant count at a feeder is a function of time and both the maximum ant count reached at a food source (Count_max_) and the time at which the ant count at a food source reaches half the maximum count (K_M_).

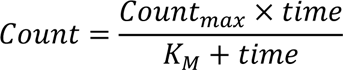

### Statistical analysis

The complete statistical analysis output, and the entire dataset on which this analysis is based, is available from Zenodo (https://doi.org/10.5281/zenodo.12073127).

All graphics and statistical analysis were generated using R version 4.2.2 (R Core Team, 2022; Wickham, 2016, 2022). Y-maze preference test proportion data was analysed using a beta regression (Cribari-Neto & Zeileis, 2010) and model fit assessed using lmtest (Zeileis & Hothorn, 2002). Individual feeding patterns and collective foraging dynamics were analysed using linear mixed-effects models (Bates *et al*., 2015) with DHARMa (Hartig, 2022) used to assess linear model assumptions and MuMIn (Bartoń, 2022) to obtain a measure of goodness of fit. Analysis of variance tables were used to test the effects of the regression’s coefficients (Fox & Weisberg, 2019). Estimated marginal means and contrasts were obtained using emmeans (Lenth, 2022) with Bonferroni adjusted values accounting for multiple testing. We avoid the use of p-values, and their associated binary decision of significant/nonsignificant, instead reporting effect size estimates and their respective 95% confidence intervals shown throughout the results section as (estimate [lower limit, upper limit, N = sample size]).

## Results

### Ants prefer the odour associated with a previously experienced food over a novel one

Colonies had no innate preference for any odour, suggesting that both apple and strawberry odour were originally equally preferred (Control: 43% of the ants preferred the odour which would later be associated with the food [35%, 51%, N = 76]; Spinosad: 54% [46%, 62%, N = 76]). However, approximately 24 hours after each colony was fed one of the four treatments, its preference for the odour of the experienced food was higher than before its ingestion (Control: 63% [55%, 71%, N = 76]; Spinosad: 67% [60%, 74%, N = 76]). Previously experiencing a food taste increased the preference for its odour by 20% [9%, 32%, N = 76] when ants were fed sucrose, and by 13% [2%, 24%, N = 76] when ants were fed spinosad-laced sucrose (Figure 2).

**Figure 2.**
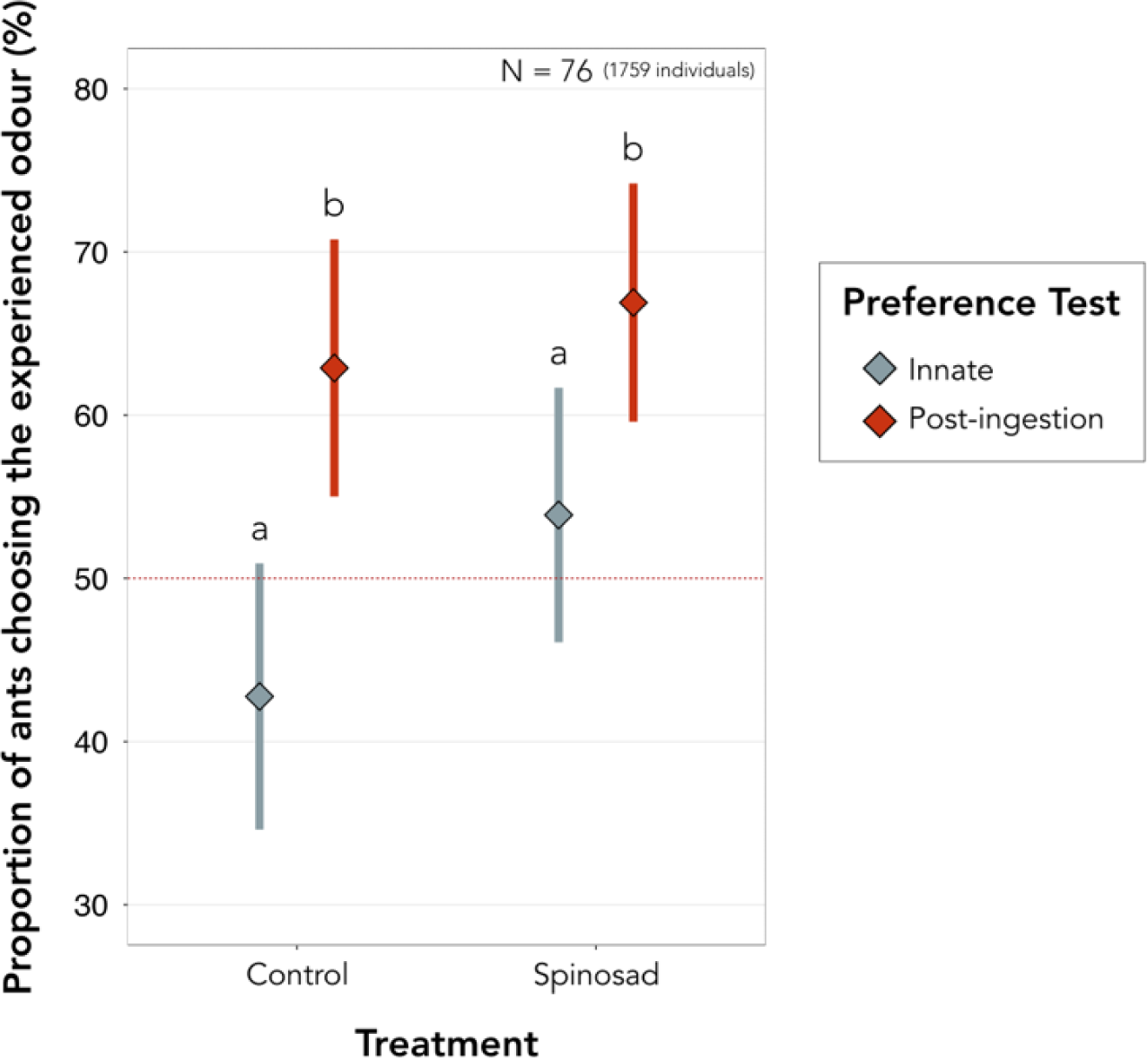
Ants prefer the odour associated with a previously (approximately 24h) experienced food over a novel one, despite having no innate preference for it. Diamonds represent the estimated marginal means obtained from the beta regression model and whiskers the respective 95% confidence intervals. Estimates of 50% (red dashed horizontal line) suggest individuals have no preference for either odour. Letters reflect statistical differences between treatments based on the estimated confidence intervals. A total of 1759 individuals from 19 colonies were tested in 76 preference tests, with each test measuring the preference of around 25 ants for either odour.

### Exposure to spinosad decreases the amount of food ingested through earlier food abandonment

Ants consumed food with a novel taste (1.9nL/s [1.6nL/s, 2.2nL/s, N = 350) faster than food with the same taste of a previously experienced food (1.7nL/s [1.4nL/s, 2.0nL/s, N = 350]), regardless of previous exposure to spinosad (Figure 3). However, exposure to spinosad led to a 24% [4%, 43% N = 350] decrease in overall food consumption relative to ants which were not exposed to the toxicant (Control: 136nL [117nL, 155nL, N = 350]; Spinosad: 112nL [95nL, 130nL, N = 350]).

**Figure 3.**
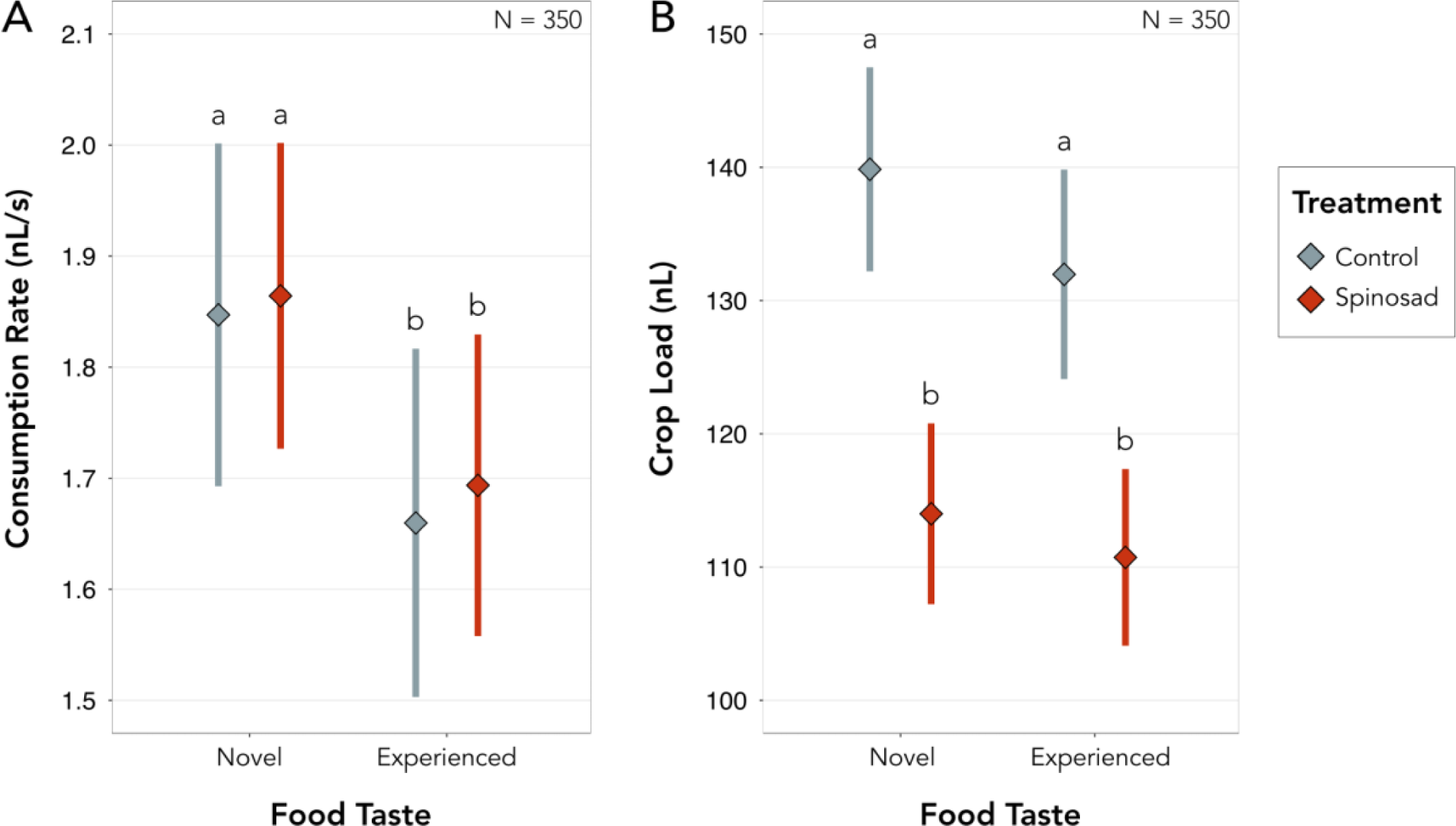
Ants consume food with a novel taste faster than that which has the same taste of a previously experienced food. Ants which ingested spinosad, a slow-acting poison, generally consume less food overall compared to those fed on the sucrose controls. Notably, as consumption rates are not affected by the presence of spinosad, decreases in crop load must result from shorter drinking events, and thus the earlier abandonment of the current food source. Diamonds represent the estimated marginal means obtained from the linear mixed-effects models and whiskers the respective standard errors. Letters reflect statistical differences between treatments based on the estimated confidence intervals.

### Ants recruit slower to a food source which tastes the same as a previously experienced food

Ant colonies recruited the same total number of ants to both available food sources, regardless of whether they were previously exposed to spinosad (Novel: 25 [16, 33, N = 38]; Experienced: 27 [18, 35, N = 38]) or simply to sucrose (Novel: 28 [19, 37, N = 38]; Experienced: 25 [16, 34, N = 38]). The recruitment dynamics towards each feeder differed with treatment (Figure 4). Under control conditions, colonies took longer to recruit towards the food which tasted the same as the previously experienced one (Novel: 6min [2min, 9min, N = 38]; Experienced: 8min [5min, 12min, N = 38]). Interestingly, this effect was lost in colonies which were previously exposed to spinosad (Novel: 4min [1min, 8min, N = 38]; Experienced: 4min [1min, 7min, N = 38]). Surprisingly, ants treated with an apple-flavoured food showed a tendency to recruit slower towards an apple tasting food, but only when it was placed on the left feeder. Moreover, the maximum number of ants recruited to a food source was affected by both odour and side, such that it reached higher values when either apple-flavoured food was placed on the right feeder or strawberry-flavoured food was placed on the left feeder. Taken together, it would appear that ants have an intrinsic preference for both strawberry-flavoured food and to turn right. Previous work has suggested *L. humile* workers have a slight preference of 58% for strawberry odour over 42% for apple (Wagner *et al*., 2023). However, previous work has also suggested these ants have a tendency to be left lateralised (Galante & Czaczkes, 2024, Poissonnier et. al. In Prep.). Given this, and since we believe such biases are satisfactorily accounted for by the full-factorial design of the experiment, we predominantly focused on the effects of pre-exposure to a food taste and to spinosad.

**Figure 4.**
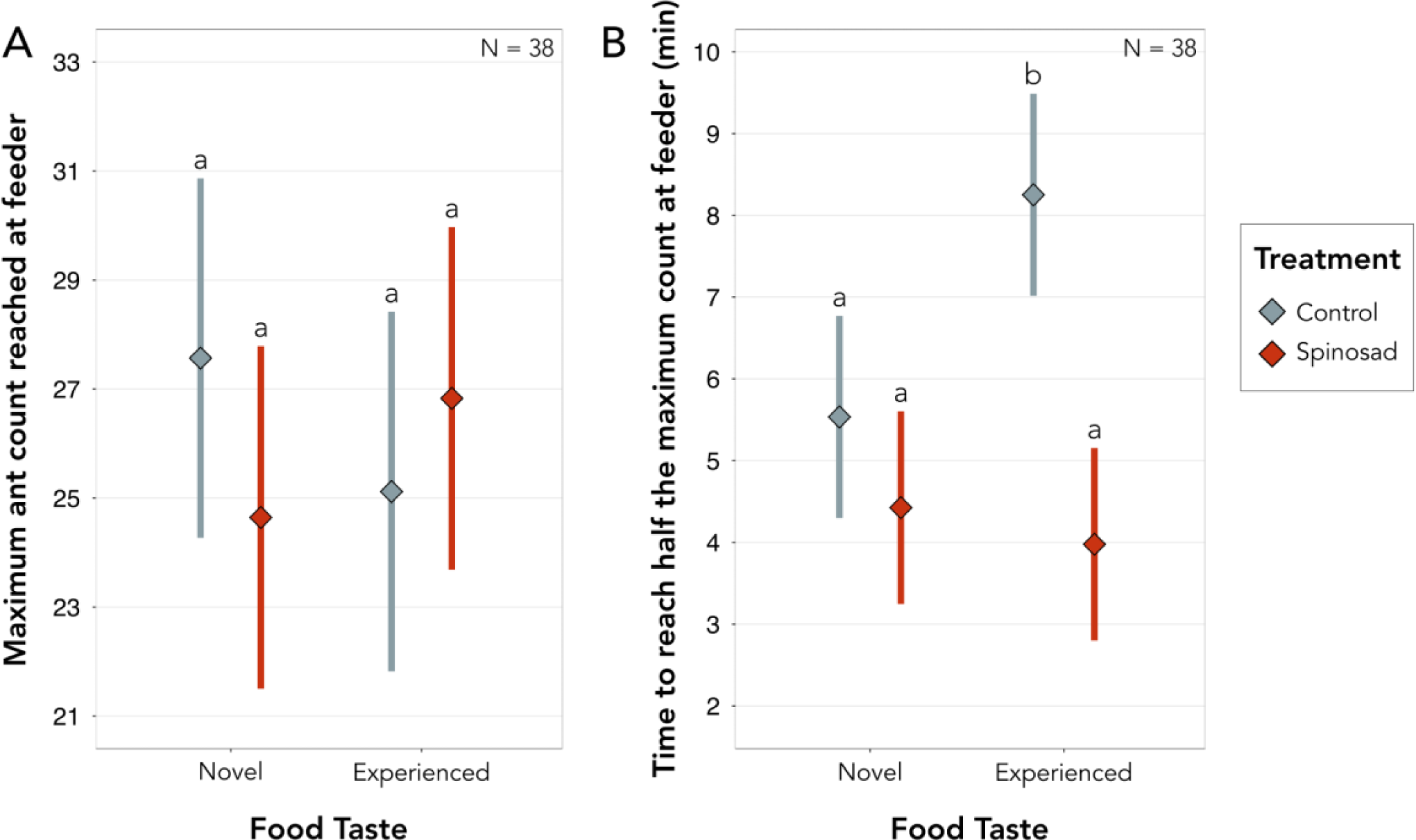
Ant colonies recruit the same number of ants to both available food sources, regardless of which food they experienced previously. However, under control conditions, colonies take longer to recruit towards a food source which tastes the same as a previously experienced one, when compared to a food source with a novel taste. Notably, this effect is lost when the previously experienced food contained spinosad, a slow-acting poison. Importantly, ants treated with an apple-flavoured food show a tendency to recruit slower towards an apple tasting food, but only when this is placed on the left feeder. Moreover, the maximum number of ants recruited to a food source was affected by both odour and side, such that it reached higher values when either apple-flavoured food was placed on the right feeder or strawberry-flavoured food was placed on the left feeder (see Statistical Analysis). Diamonds represent the estimated marginal means obtained from the linear mixed-effects models and whiskers the respective standard errors. Letters reflect statistical differences between treatments based on the estimated confidence intervals.

## Discussion

Ants prefer the odour of a known food, over that of a novel food, regardless of spinosad exposure (Figure 2). Previous studies have also shown that ants strongly prefer the first odour associated with food which they experience after food deprivation (Oberhauser, Bogenberger & Czaczkes, 2022). Had ants been capable of forming a conditioned taste aversion (CTA) to spinosad, they would have been expected to actively prefer a novel food taste over a previously experienced one when this was paired with the toxicant. The lack of aversion observed could be due to the sublethal dose of spinosad used not causing post-ingestion malaise. In fact, during the experiment we observed a general lack of mortality, which we anecdotally quantified. In control-treated colonies, we observed a worker mortality of 5.4% ± 1.9% (N = 3), with a mortality of marked individuals which were directly fed the spinosad-laced solution of 7.3% ± 6.5% (N = 3). Similarly, spinosad-treated colonies had a mortality of 8.3% ± 2.3% (N = 5), with a mortality of marked individuals which were directly fed the treatment of 7.0% ± 2.5% (N = 5). This represents a spinosad associated mortality of around 3%, which was unexpected as previous work suggested Argentine ants, in groups of four individuals, fed a similar dose of spinosad had a mortality likelihood of 19% 24 hours post-ingestion. This is likely a consequence of group size, such that mortality is higher in small groups than in larger groups for the same toxicant dose (Galante et. al., In Prep.).

Spinosad-laced sucrose solutions were readily accepted by the ants, likely because they either could not detect the presence of the toxicant or did not recognise it as toxic on a first contact. However, even at the extremely small dose of spinosad used, which is around 600 times less than that typically used in field conditions (Milosavljević *et al*., 2024, Pedraza et. al., In prep.), spinosad had a clear post-ingestion effect: individuals exposed to the toxicant showed a decrease in overall consumption of roughly 20% on all foods ingested afterwards, even though none of these foods contained spinosad. This suggests a single exposure to spinosad makes ants ingest less food for at least a day post-exposure. This may be driven by either physiological or cognitive processes. Moreover, while ants ingested food with a novel taste at a faster rate than food with a familiar taste, consumption rate was not affected by exposure to spinosad (Figure 3). Thus, the observed decrease in crop load must have been a result of shorter drinking events and the earlier abandonment of the food source. However, trail pheromones have been found to alter Argentine ant’s subjective evaluation of food, increasing their acceptance (Rossi *et al*., 2020). Thus, during collective foraging, the decrease in food consumption found at the individual level might be mitigated by the presence of trail pheromone.

CTA is often characterised by a single conditioning trial with a long temporal separation between food ingestion and post-ingestion malaise (Nachman, 1970; Steinert, Infurna & Spear, 1980; Rosas & Bouton, 1996). However, crickets have been shown to successfully form associations only when the two stimuli were separated by less than one hour (Lyu & Mizunami, 2022). In our case, the sublethal dose of spinosad used might have taken too long to cause post-ingestion malaise, which could have prevented the formation of CTA. As a reference, a field-realistic concentration of 150ppm spinosad showed a survival likelihood of 72% after one hour whilst 0.25ppm resulted in a survival likelihood of 81% after 24 hours (Galante et. al., In Prep.). With that said, the level of spinosad used did cause a reduction in feeding of around 20%, which would be consistent with ongoing malaise.

Under the control conditions, colonies took longer to recruit to previously experienced food sources than to novel ones (Figure 4). This was likely due to ants which foraged on a novel food source ingesting it faster, and thus return to the nest sooner, than those feeding on a previously experienced food taste. Mass-recruiting ants, such as *L. humile*, rely on pheromone trails during foraging, with ants laying more pheromone towards higher quality food sources (Latty *et al*., 2017). Small initial differences between pheromone trails often result in large recruitment differences and in most cases in preference for one food source over the other, a phenomenon often called symmetry breaking (Beckers *et al*., 1990; Sumpter & Beekman, 2003; Detrain & Deneubourg, 2008; Grüter *et al*., 2012). Interestingly, despite initial differences in recruitment, control-treated colonies reach the same number of individuals at both food sources and no symmetry breaking was observed. This lack of preference for one food source over the other is thought to be one way by which small colonies exploit multiple food sources in order to avoid large-scale conflicts (Nicolis & Deneubourg, 1999) which would be too costly since the monopolisation of resources through cooperative defence cannot be efficiently implemented by small colonies (Franks & Partridge, 1993; Mailleux, Deneubourg & Detrain, 2003). Moreover, the lack of symmetry breaking observed is likely a consequence of both food sources being of equally high energetic value (Price *et al*., 2016) and due to the negative feedback generated by overcrowding at a food source (Czaczkes, Grüter & Ratnieks, 2013; Wendt, Kleinhoelting & Czaczkes, 2020). The lack of brood in the colony is also known to decrease collective foraging asymmetry (Portha, 2002) and this was also likely potentiated by the two food sources being in close proximity to each other (13cm apart and 15cm from the nest entrances).

The initial recruitment differences observed in control-treated colonies were not present in spinosad-treated ones. Spinosad exposure resulted in equally fast recruitment for food sources of both novel and experienced taste, and even slightly faster recruitment than control colonies overall (Figure 4). However, individual level consumption rates did not differ with exposure to spinosad. Colonies feeding on a novel food source would still ingest it faster, and thus return to the nest sooner, than those feeding on a previously experienced food taste. A potential explanation for this would be that exposure to spinosad results in weaker pheromone trail deposition or affects the ant’s ability to detect pheromone trails. Ants are capable of regulating pheromone deposition (Hölldobler, Stanton & Markl, 1978; Beckers, Deneubourg & Goss, 1992; Jackson & Châline, 2007), and exposure to spinosad resulted in reduced consumption of all food sources. When pheromone trail intensity is low, the decision of individual ants to deposit pheromone will have a disproportionate effect on the relative trail strength (Price *et al*., 2016). Moreover, weaker pheromone trails could lead ants to rely more on private information, such as learnt memories. Argentine ants have been shown to predominantly rely on social cues, such as pheromone trails, more heavily than on private information (Aron *et al*., 1993; Von Thienen *et al*., 2014; Von Thienen, Metzler & Witte, 2016). However, other mass-recruiting ants have been shown to combine both information sources for more efficient foraging or even to rely more heavily on route memories, which are often more accurate than pheromone trails, when these conflict with social cues (Evison *et al*., 2008; Grüter, Czaczkes & Ratnieks, 2011; Czaczkes *et al*., 2011). Similarly, Argentine ants have been shown to choose between trails randomly when pheromone concentration is low (Von Thienen *et al*., 2014).

Alternatively, spinosad could induce a state of hyperactivity, causing ants to move faster, be more motivated to find food or even forage more efficiently. In fact, Argentine ants fed moderate doses of caffeine, a neuroactive chemical, have been shown to have improved foraging (Galante *et al*., 2024). Spinosad, predominantly acting on nicotinic acetylcholine receptors, has also been identified as an antagonist of GABA receptors (see Millar & Denholm, 2007 and Kirst, 2010 for reviews). GABA, the main inhibitory neurotransmitter in invertebrates (Nepi, 2014) is often linked to an animal’s ability to forget, and thus adapt to its environment (Boitard *et al*., 2015). By preventing the release of GABA, spinosad could lead ants to fixate on initially appetitive spinosad-laced sucrose solutions, potentially explaining the observed improvements in recruitment from spinosad exposed colonies. Nevertheless, a recent study showed GABA promoting flower fidelity, suggesting its effects are complex (Calderai *et al*., 2023). Interestingly, sub-lethal doses of the neonicotinoid imidacloprid, also an antagonist of nicotinic acetylcholine receptors, were shown to shift colony-level preference in the invasive ant *Lasius neglectus* towards toxicant-laced solutions (Frizzi *et al*., 2022), even though imidacloprid has been shown to impair olfactory learning and memory in honeybees (Yang *et al*., 2012; Williamson & Wright, 2013). However, were this the case, control attempts using spinosad as the toxicant would likely have been more successful, unless such effects are lost at high doses, as they are for caffeine and imidacloprid (Frizzi *et al*., 2022; Galante *et al*., 2024).

The sublethal dose of spinosad used in our experiment did not lead the ants to form a conditioned taste aversion, potentially because of its low toxicity and the high energy content of the sucrose solutions provided. Nevertheless, conditioned taste aversions could still be present under field-conditions, where toxicant doses are higher. If this is the case, a potential way to address CTA in management efforts would be to leverage these aversive associations. For example, using toxic baits with one taste-odour initially, and once individuals formed a CTA to that taste-odour, cycle through different flavours such that individuals would always be driven towards novel foods, yet ingestion of the toxicant would remain. However, this approach would need to be formally tested as natural food sources would still be competing with the baits. Ultimately, identifying a toxicant which is not only palatable but also has no direct impact on food consumption is crucial for the development of effective control strategies.

## Acknowledgements

We thank E. Sequeira and S. Abril for ant collection, and LA. Poissonnier for helpful discussions and feedback.

## Ethical statement

We have conducted all experiments in accordance with the guidelines that are applicable to working with the model organism in the European Union. Colonies were kept in closed boxes under oil baths in order to prevent any escape.

## Author contributions

**H. Galante** Conceptualization, Methodology, Software, Validation, Formal analysis, Investigation, Data Curation, Writing - Original Draft, Writing - Review & Editing, Visualization, Supervision. **M. Forster:** Investigation, Writing - Review & Editing. **C. Werneke:** Formal analysis, Writing - Review & Editing. **T. J. Czaczkes:** Conceptualization, Methodology, Resources, Writing - Review & Editing, Supervision, Project administration, Funding acquisition.

## References

Angulo, E., Guénard, B., Balzani, P., Bang, A., et al. (2024) The Argentine ant, *Linepithema humile*: natural history, ecology and impact of a successful invader. Entomologia Generalis. 44 (1), 41–61. 10.1127/entomologia/2023/2187.

Angulo, E., Hoffmann, B.D., Ballesteros-Mejia, L., Taheri, A., et al. (2022) Economic costs of invasive alien ants worldwide. Biological Invasions. 24, 2041–2060. 10.1007/s10530-022-02791-w.

Arenas, A. & Roces, F. (2018) Appetitive and aversive learning of plants odors inside different nest compartments by foraging leaf-cutting ants. Journal of Insect Physiology. 109, 85–92. 10.1016/j.jinsphys.2018.07.001.

Aron, S., Beckers, R., Deneubourg, J.L. & Pasteels, J.M. (1993) Memory and chemical communication in the orientation of two mass-recruiting ant species. Insectes Sociaux. 40 (4), 369–380. 10.1007/BF01253900.

Ayestaran, A., Giurfa, M. & Sanchez, M.G.B. (2010) Toxic but drank: gustatory aversive compounds induce post-ingestional malaise in harnessed honeybees. PLoS ONE. 5 (10), e15000. 10.1371/journal.pone.0015000.

Babin, A., Kolly, S., Schneider, F., Dolivo, V., et al. (2014) Fruit flies learn to avoid odours associated with virulent infection. Biology Letters. 10 (3), 20140048. 10.1098/rsbl.2014.0048.

Bacci, L., Lupi, D., Savoldelli, S. & Rossaro, B. (2016) A review of Spinosyns, a derivative of biological acting substances as a class of insecticides with a broad range of action against many insect pests. Journal of Entomological and Acarological Research. 48 (1), 40. 10.4081/jear.2016.5653.

Bartoń, K. (2022) MuMIn: Multi-model inference. https://CRAN.R-project.org/package=MuMIn.

Bates, D., Mächler, M., Bolker, B. & Walker, S. (2015) Fitting linear mixed-effects models using lme4. Journal of Statistical Software. 67 (1), 1–48. 10.18637/jss.v067.i01.

Beckers, R., Deneubourg, J. & Goss, S. (1992) Trail laying behaviour during food recruitment in the ant *Lasius niger* (L.). Insectes Sociaux. 39 (1), 59–72. 10.1007/BF01240531.

Beckers, R., Deneubourg, J.L., Goss, S. & Pasteels, J.M. (1990) Collective decision making through food recruitment. Insectes Sociaux. 37 (3), 258–267. 10.1007/BF02224053.

Bernays, E.A. & Lee, J.C. (1988) Food aversion learning in the polyphagous grasshopper *Schistocerca americana*. Physiological Entomology. 13 (2), 131–137. 10.1111/j.1365-3032.1988.tb00916.x.

Bernstein, I.L. (1999) Taste aversion learning: a contemporary perspective. Nutrition. 15 (3), 229–234. 10.1016/S0899-9007(98)00192-0.

Biondi, A., Mommaerts, V., Smagghe, G., Viñuela, E., et al. (2012) The non-target impact of spinosyns on beneficial arthropods. Pest Management Science. 68 (12), 1523–1536. 10.1002/ps.3396.

Boitard, C., Devaud, J., Isabel, G. & Giurfa, M. (2015) GABAergic feedback signaling into the calyces of the mushroom bodies enables olfactory reversal learning in honey bees. Frontiers in Behavioral Neuroscience. 9. 10.3389/fnbeh.2015.00198.

Bruin, M.K., Röst, L.C.M. & Draisma, F.G.A.M. (1977) Estimates of the number of foraging ants with the Lincoln-Index method in relation to the colony size of *Formica polyctena*. Journal of Animal Ecology. 46 (2), 457–470. 10.2307/3823.

Cabrera, E., Fontan, I.R., Hoffmann, B.D. & Josens, R. (2021) Laboratory and field insights into the dynamics and behavior of Argentine ants, *Linepithema humile*, feeding from hydrogels. Pest Management Science. 77 (7), 3250–3258. 10.1002/ps.6368.

Calderai, G., Baggiani, B., Bianchi, S., Pierucci, V., et al. (2023) Nectar-borne GABA promotes flower fidelity in bumble bees. Entomologia Generalis. 10.1127/entomologia/2023/2062.

Cribari-Neto, F. & Zeileis, A. (2010) Beta Regression in R. Journal of Statistical Software. 34 (2), 1–24. 10.18637/jss.v034.i02.

Czaczkes, T.J. (2018) Using T- and Y-mazes in myrmecology and elsewhere: A practical guide. Insectes Sociaux. 65 (2), 213–224. 10.1007/s00040-018-0621-z.

Czaczkes, T.J., Grüter, C., Jones, S.M. & Ratnieks, F.L.W. (2011) Synergy between social and private information increases foraging efficiency in ants. Biology Letters. 7 (4).

Czaczkes, T.J., Grüter, C. & Ratnieks, F.L.W. (2013) Negative feedback in ants: crowding results in less trail pheromone deposition. Journal of The Royal Society Interface. 10 (81). 10.1098/rsif.2012.1009.

Czaczkes, T.J. & Kumar, P. (2020) Very rapid multi-odour discrimination learning in the ant *Lasius niger*. Insectes Sociaux. 67 (4), 541–545. 10.1007/s00040-020-00787-0.

Desmedt, L., Baracchi, D., Devaud, J., Giurfa, M., et al. (2017) Aversive learning of odor-heat associations in ants. Journal of Experimental Biology. jeb.161737. 10.1242/jeb.161737.

Dethier, V.G. (1980) Food-aversion learning in two polyphagous caterpillars, *Diacrisia virginica* and *Estigmene congrua*. Physiological Entomology. 5 (4), 321–325. 10.1111/j.1365-3032.1980.tb00242.x.

Detrain, C. & Deneubourg, J. (2008) Collective decision-making and foraging patterns in ants and honeybees. Advances in Insect Physiology. 35, 123–173. 10.1016/S0065-2806(08)00002-7.

Dickinson, A. (2012) Associative learning and animal cognition. Philosophical Transactions of the Royal Society B: Biological Sciences. 367 (1603), 2733–2742. 10.1098/rstb.2012.0220.

Evison, S.E.F., Petchey, O.L., Beckerman, A.P. & Ratnieks, F.L.W. (2008) Combined use of pheromone trails and visual landmarks by the common garden ant *Lasius niger*. Behavioral Ecology and Sociobiology. 63 (2), 261–267. 10.1007/s00265-008-0657-6.

Fox, J. & Weisberg, S. (2019) An R companion to applied regression. Third. Thousand Oaks CA, Sage. https://socialsciences.mcmaster.ca/jfox/Books/Companion/.

Franks, N.R. & Partridge, L. (1993) Lanchester battles and the evolution of combat in ants. Animal Behaviour. 45 (1), 197–199. https://psycnet.apa.org/doi/10.1006/anbe.1993.1021.

Frizzi, F., Balzani, P., Masoni, A., Wendt, C.F., et al. (2022) Sub-lethal doses of imidacloprid alter food selection in the invasive garden ant *Lasius neglectus*. Environmental Science and Pollution Research. 10.1007/s11356-022-24100-7.

Galante, H. & Czaczkes, T.J. (2024) Invasive ant learning is not affected by seven potential neuroactive chemicals. Current Zoology. 70 (1), 87–97. 10.1093/cz/zoad001.

Galante, H., Czaczkes, T.J. & De Agrò, M. (2024) Three-dimensional body reconstruction enables quantification of liquid consumption in small invertebrates. bioRxiv. 10.1101/2024.06.14.599002.

Galante, H., De Agrò, M., Koch, A., Kau, S., et al. (2024) Acute exposure to caffeine improves foraging in an invasive ant. iScience. 10.1016/j.isci.2024.109935.

Garcia, J. & Koelling, R.A. (1966) Relation of cue to consequence in avoidance learning. Psychonomic Science. 4 (1), 123–124. 10.3758/BF03342209.

Grüter, C., Czaczkes, T.J. & Ratnieks, F.L.W. (2011) Decision making in ant foragers (*Lasius niger*) facing conflicting private and social information. Behavioral Ecology and Sociobiology. 65 (2), 141–148. 10.1007/s00265-010-1020-2.

Grüter, C., Schürch, R., Czaczkes, T.J., Taylor, K., et al. (2012) Negative feedback enables fast and flexible collective decision-making in ants. PLoS ONE. 7 (9), e44501. 10.1371/journal.pone.0044501.

Hartig, F. (2022) DHARMa: Residual diagnostics for hierarchical (multi-level/mixed) regression models. https://CRAN.R-project.org/package=DHARMa.

Hölldobler, B., Stanton, R.C. & Markl, H. (1978) Recruitment and food-retrieving behavior in *Novomessor* (Formicidae, Hymenoptera). Behavioral Ecology and Sociobiology. 4, 163–181. 10.1007/BF00354978.

Hölldobler, B. & Wilson, E.O. (2009) The superorganism: the beauty elegance and strangeness of insect societies. WW Norton & Company.

Jackson, D.E. & Châline, N. (2007) Modulation of pheromone trail strength with food quality in Pharaoh’s ant, *Monomorium pharaonis*. Animal Behaviour. 74 (3), 463–470. 10.1016/j.anbehav.2006.11.027.

Khan, H.A.A. (2018) Spinosad resistance affects biological parameters of *Musca domestica* Linnaeus. Scientific Reports. 8 (1), 14031. 10.1038/s41598-018-32445-8.

Kirst, H.A. (2010) The spinosyn family of insecticides: realizing the potential of natural products research. The Journal of Antibiotics. 63 (3), 101–111. 10.1038/ja.2010.5.

Knaden, M. & Graham, P. (2016) The sensory ecology of ant navigation: From natural environments to neural mechanisms. Annual Review of Entomology. 61 (1), 63–76. 10.1146/annurev-ento-010715-023703.

Kobler, J.M., Jimenez, F.J.R., Petcu, I. & Kadow, I.C.G. (2020) Immune receptor signaling and the mushroom body mediate post-ingestion pathogen avoidance. Current Biology. 30 (23), 4693–4709. 10.1016/j.cub.2020.09.022.

Lai, Y., Despouy, E., Sandoz, J., Su, S., et al. (2020) Degradation of an appetitive olfactory memory via devaluation of sugar reward is mediated by 5-HT signaling in the honey bee. Neurobiology of Learning and Memory. 173, 107278. 10.1016/j.nlm.2020.107278.

Latty, T., Holmes, M.J., Makinson, J.C. & Beekman, M. (2017) Argentine ants (*Linepithema humile*) use adaptable transportation networks to track changes in resource quality. Journal of Experimental Biology. 220 (4), 686–694. 10.1242/jeb.144238.

Lenth, R.V. (2022) emmeans: Estimated marginal means, aka least-squares means. https://CRAN.R-project.org/package=emmeans.

Lewis, T., Pollard, G.V. & Dibley, G.C. (1974) Rhythmic foraging in the leaf-cutting ant *Atta cephalotes* (L.) (Formicidae: Attini). Journal of Animal Ecology. 43 (1), 129–141. 10.2307/3162.

Lyu, H. & Mizunami, M. (2022) Conditioned taste aversion in the cricket *Gryllus bimaculatus*. Scientific Reports. 12 (1), 9751. 10.1038/s41598-022-13500-x.

Mailleux, A., Deneubourg, J. & Detrain, C. (2003) How does colony growth influence communication in ants? Insectes Sociaux. 50 (1), 24–31. 10.1007/s000400300004.

Mathis, A., Mamidanna, P., Cury, K.M., Abe, T., et al. (2018) DeepLabCut: Markerless pose estimation of user-defined body parts with deep learning. Nature Neuroscience. 21, 1281–1289. 10.1038/s41593-018-0209-y.

Michaelis, L. & Menten, M.L. (1913) Die kinetik der invertinwirkung. Biochemische Zeitschrift. 49, 333–369. https://www.chem.uwec.edu/Chem352_Resources/pages/readings/media/Michaelis_&_Menton_1913.pdf.

Millar, N.S. & Denholm, I. (2007) Nicotinic acetylcholine receptors: targets for commercially important insecticides. Invertebrate Neuroscience. 7 (1), 53–66. 10.1007/s10158-006-0040-0.

Milosavljević, I., Irvin, N.A., Lewis, M. & Hoddle, M.S. (2024) Spinosad-infused biodegradable hydrogel beads as a potential organic approach for argentine ant, *Linepithema humile* (Mayr) (Hymenoptera: Formicidae), management in California citrus orchards. Journal of Applied Entomology. 148 (1), 117–127. 10.1111/jen.13203.

Nachman, M. (1970) Learned taste and temperature aversions due to lithium chloride sickness after temporal delays. Journal of Comparative and Physiological Psychology. 73 (1), 22–30. 10.1037/h0029807.

Nakai, J., Totani, Y., Hatakeyama, D., Dyakonova, V.E., et al. (2020) Another example of conditioned taste aversion: case of snails. Biology. 9 (12), 422. 10.3390/biology9120422.

Nath, T., Mathis, A., Chen, A.C., Patel, A., et al. (2019) Using DeepLabCut for 3D markerless pose estimation across species and behaviors. Nature Protocols. 14, 2152–2176. 10.1038/s41596-019-0176-0.

Nepi, M. (2014) Beyond nectar sweetness: The hidden ecological role of non-protein amino acids in nectar. Journal of Ecology. 102 (1), 108–115. 10.1111/1365-2745.12170.

Nicolis, S.C. & Deneubourg, J.L. (1999) Emerging patterns and food recruitment in ants: an analytical study. Journal of Theoretical Biology. 198 (4), 575–592. 10.1006/jtbi.1999.0934.

Oberhauser, F.B., Bogenberger, K. & Czaczkes, T.J. (2022) Ants prefer the option they are trained to first. Journal of Experimental Biology. 225 (24), jeb243984. 10.1242/jeb.243984.

Oberhauser, F.B., Schlemm, A., Wendt, S. & Czaczkes, T.J. (2019) Private information conflict: *Lasius niger* ants prefer olfactory cues to route memory. Animal Cognition. 22 (3), 355–364. 10.1007/s10071-019-01248-3.

Palmerino, C.C., Rusiniak, K.W. & Garcia, J. (1980) Flavor-illness aversions: the peculiar roles of odor and taste in memory for poison. Science. 208 (4445), 753–755. 10.1126/science.7367891.

Pavlov, I.P. (1927) *Conditioned reflexes: An investigation of the physiological activity of the cerebral cortex*. Oxford, England, Oxford University Press. https://psycnet.apa.org/record/1927-02531-000.

Piqueret, B., Sandoz, J. & d’Ettorre, P. (2019) Ants learn fast and do not forget: Associative olfactory learning, memory and extinction in *Formica fusca*. Royal Society Open Science. 6 (6). 10.1098/rsos.190778.

Porter, S.D. & Jorgensen, C.D. (1981) Foragers of the harvester ant, *Pogonomyrmex owyheei*: a disposable caste? Behavioral Ecology and Sociobiology. 9, 247–256. 10.1007/BF00299879.

Portha, S. (2002) Self-organized asymmetries in ant foraging: a functional response to food type and colony needs. Behavioral Ecology. 13 (6), 776–781. 10.1093/beheco/13.6.776.

Price, R.I., Grüter, C., Hughes, W.O.H. & Evison, S.E.F. (2016) Symmetry breaking in mass-recruiting ants: extent of foraging biases depends on resource quality. Behavioral Ecology and Sociobiology. 70 (11), 1813–1820. 10.1007/s00265-016-2187-y.

R Core Team (2022) *R: A language and environment for statistical computing*. Vienna, Austria, R Foundation for Statistical Computing. https://www.R-project.org/.

Rescorla, R.A. & Wagner, A.R. (1972) A theory of Pavlovian conditioning: Variations in the effectiveness of reinforcement and nonreinforcement. In: Classical conditioning II: Current research and theory. Black, A. H., Prokasy, W. F. New York, Appleton-Century-Crofts. pp. 64–99.

Rosas, J.M. & Bouton, M.E. (1996) Spontaneous recovery after extinction of a conditioned taste aversion. Animal Learning & Behavior. 24 (3), 341–348. 10.3758/BF03198982.

Rossi, N., Pereyra, M., Moauro, M.A., Giurfa, M., et al. (2020) Trail-pheromone modulates subjective reward evaluation in Argentine ants. Journal of Experimental Biology. 223 (17). 10.1242/jeb.230532.

Salgado, V.L. (1998) Studies on the mode of action of spinosad: insect symptoms and physiological correlates. Pesticide Biochemistry and Physiology. 60 (2), 91–102. 10.1006/pest.1998.2332.

Schindelin, J., Arganda-Carreras, I., Frise, E., Kaynig, V., et al. (2012) Fiji: an open-source platform for biological-image analysis. Nature Methods. 9 (7), 676–682. 10.1038/nmeth.2019.

Schneider, C.A., Rasband, W.S. & Eliceiri, K.W. (2012) NIH Image to ImageJ: 25 years of image analysis. Nature Methods. 9, 671–675. 10.1038/nmeth.2089.

Silverman, J. & Brightwell, R.J. (2008) The Argentine ant: challenges in managing an invasive unicolonial pest. Annual Review of Entomology. 53 (1), 231–252. 10.1146/annurev.ento.53.103106.093450.

Simões, P.M.V., Ott, S.R. & Niven, J.E. (2012) A long-latency aversive learning mechanism enables locusts to avoid odours associated with the consequences of ingesting toxic food. Journal of Experimental Biology. 215 (10), 1711–1719. 10.1242/jeb.068106.

Steinert, P.A., Infurna, R.N. & Spear, N.E. (1980) Long-term retention of a conditioned taste aversion in preweanling and adult rats. Animal Learning & Behavior. 8 (3), 375–381. 10.3758/BF03199620.

Sumpter, D.J.T. & Beekman, M. (2003) From nonlinearity to optimality: pheromone trail foraging by ants. Animal Behaviour. 66 (2), 273–280. 10.1006/anbe.2003.2224.

Varnon, C.A., Dinges, C.W., Black, T.E., Wells, H., et al. (2018) Failure to find ethanol-induced conditioned taste aversion in honey bees (*Apis mellifera* L.). Alcoholism: Clinical and Experimental Research. 42 (7), 1260–1270. 10.1111/acer.13761.

Von Thienen, W., Metzler, D., Choe, D. & Witte, V. (2014) Pheromone communication in ants: a detailed analysis of concentration-dependent decisions in three species. Behavioral Ecology and Sociobiology. 68 (10), 1611–1627. 10.1007/s00265-014-1770-3.

Von Thienen, W., Metzler, D. & Witte, V. (2016) How memory and motivation modulate the responses to trail pheromones in three ant species. Behavioral Ecology and Sociobiology. 70 (3), 393–407. 10.1007/s00265-016-2059-5.

Wagner, T., Galante, H., Josens, R. & Czaczkes, T.J. (2023) Systematic examination of learning in the invasive ant *Linepithema humile* reveals fast learning and long-lasting memory. Animal Behaviour. 203, 41–52. 10.1016/j.anbehav.2023.06.012.

Wendt, S., Kleinhoelting, N. & Czaczkes, T.J. (2020) Negative feedback: ants choose unoccupied over occupied food sources and lay more pheromone to them. Journal of The Royal Society Interface. 17 (163), 20190661. 10.1098/rsif.2019.0661.

Wenig, K., Bach, R. & Czaczkes, T.J. (2021) Hard limits to cognitive flexibility: ants can learn to ignore but not avoid pheromone trails. Journal of Experimental Biology. 224 (11). 10.1242/jeb.242454.

Wickham, H. (2016) *ggplot2: Elegant graphics for data analysis*. Springer-Verlag New York. https://ggplot2.tidyverse.org.

Wickham, H. (2022) stringr: Simple, consistent wrappers for common string operations. R package version 1.5.0. https://CRAN.R-project.org/package=stringr.

Williamson, S.M. & Wright, G.A. (2013) Exposure to multiple cholinergic pesticides impairs olfactory learning and memory in honeybees. Journal of Experimental Biology. jeb.083931. 10.1242/jeb.083931.

Wright, G. (2011) The role of dopamine and serotonin in conditioned food aversion learning in the honeybee. Communicative & Integrative Biology. 4 (3), 318–320. 10.4161/cib.4.3.14840.

Wright, G.A., Mustard, J.A., Simcock, N.K., Ross-Taylor, A.A.R., et al. (2010) Parallel reinforcement pathways for conditioned food aversions in the honeybee. Current Biology. 20 (24), 2234–2240. 10.1016/j.cub.2010.11.040.

Yang, E., Chang, H., Wu, W. & Chen, Y. (2012) Impaired olfactory associative behavior of honeybee workers due to contamination of imidacloprid in the larval stage. PLoS ONE. 7 (11), e49472. 10.1371/journal.pone.0049472.

Yen, J., Chang, F. & Chang, S. (1995) A new criterion for automatic multilevel thresholding. IEEE Transactions on Image Processing. 4 (3), 370–378. 10.1109/83.366472.

Zanola, D., Czaczkes, T.J. & Josens, R. (2024) Ants evade harmful food by active abandonment. Communications Biology. 7 (1), 84. 10.1038/s42003-023-05729-7.

Zeileis, A. & Hothorn, T. (2002) Diagnostic checking in regression relationships. R News. 2 (3), 7–10. https://CRAN.R-project.org/doc/Rnews/.

